## Supplementary figures for "Membrane binding controls ordered self-assembly of animal septins"

### Figure supplements

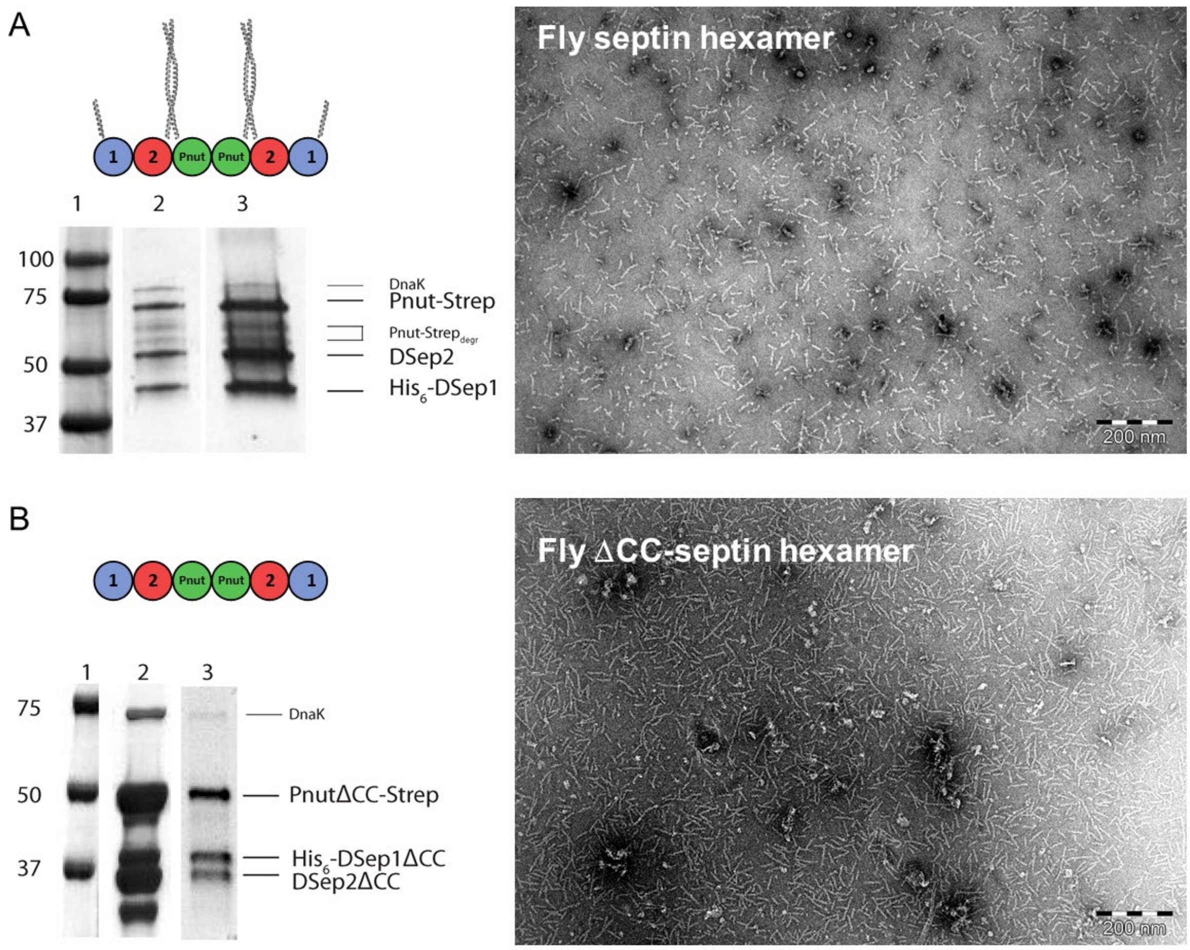


**Figure 1 – Figure supplement 1. Biochemical and morphological characterization of septins.** (A) Left: SDS-PAGE gel of full-length fly septin hexamers with lanes showing: 1) reference marker (molecular weights in kDa indicated on the left), 2) nickel column eluate, 3) StrepTrap-column eluate. The three bands of similar intensity represent the three subunits: Pnut-Strep (61.2 kDa), DSep2 (48.5 kDa) and His$_6$-DSep1 (43.8 kDa). The fainter bands between the DSep2 and Pnut bands represent partially degraded Pnut-Strep, while the faint band located around 70 kDa is DnaK [1]. Right: negative-stain TEM image showing that fly septins in a high-salt (300 mM) storage buffer are present predominantly as hexamers or dimers thereof. (B) Left: SDS-PAGE gel of the ΔCC mutant with C-terminal truncations (DSep1ΔC56, DSep2ΔC111, PnutΔC123), showing 3 bands at the positions expected for the three septin subunits after the nickel (lane 2) and StrepTrap-column (lane 3). Right: negative-stain TEM image shows that the ΔCC-septins in a high-salt (300 mM) storage buffer are present mainly as hexamers.

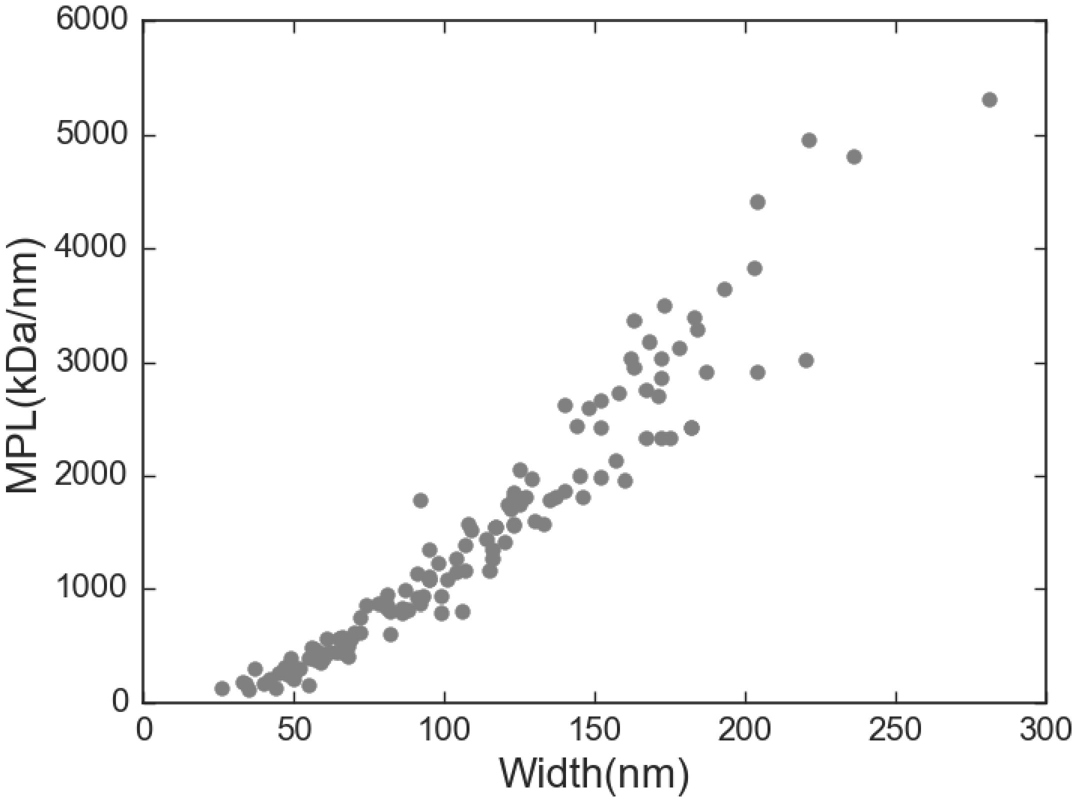


**Figure 1 – Figure supplement 2. Quantification of the bundle size for septin bundles formed in bulk solution at 500 nM.** Scatter plot of the MPL (in units of kDa/nm) versus the bundle width (in nm). The data were collected from 2 independent STEM-imaging sessions, 5 images, 16 bundles, and in total 130 data points. The dotted lines show a linear dependence expected in case of hollow cylinders or flat filament arrays and a quadratic dependence expected for solid cylinders (see legend). The lines are based on fits to small-width (<100 nm) bundles, as in this regime there was the highest number of data points.

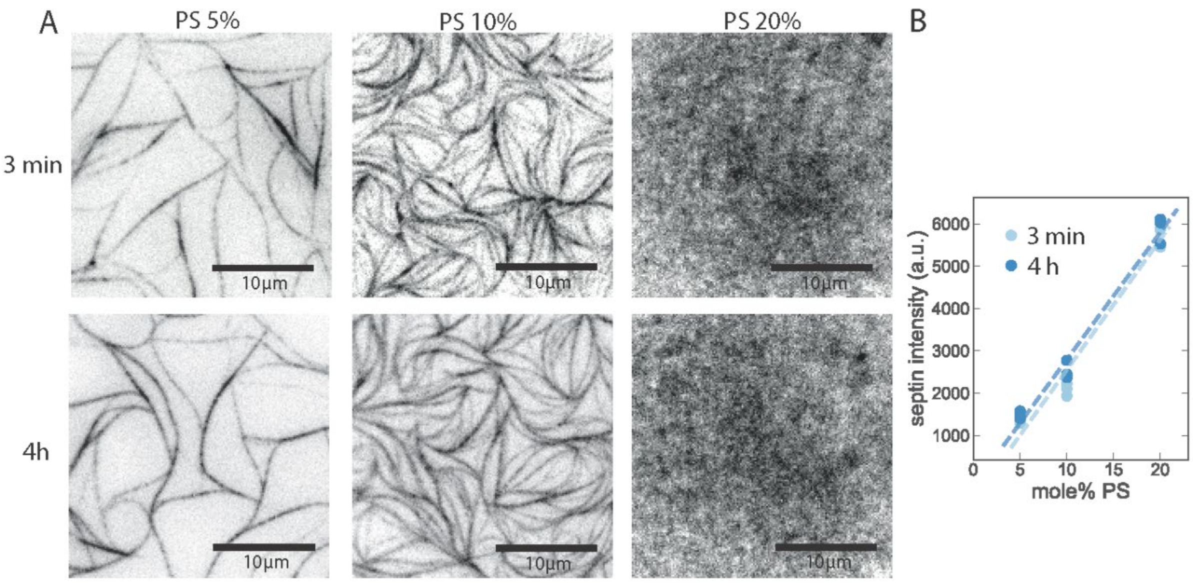


**Figure 2 – Figure supplement 1: TIRF imaging shows that fly septin hexamers rapidly adsorb to anionic bilayers and rapidly form filaments.** (A) TIRF images obtained at time points of ~3 min (top row) and 4 h (bottom row) after the deposition of mEGFP-tagged fly septins (500 nM) on PC bilayers doped with 5, 10 or 20 mol-% PS. The larger the DOPS content, the less bundled and the more dense and grainy the septin layer appears. Note that the 3 min and 4 h images do not show the same regions because experiments required keeping the glass slides in a humidified petri dish, so finding the same region afterwards was not possible. (B) Corresponding integrated septin intensities (each data point is based on one experiment with 3 different regions of interest in different sample locations), showing a linear increase with PS content (lines are fits) with insignificant changes from 3 min to 4 h of septin incubation.

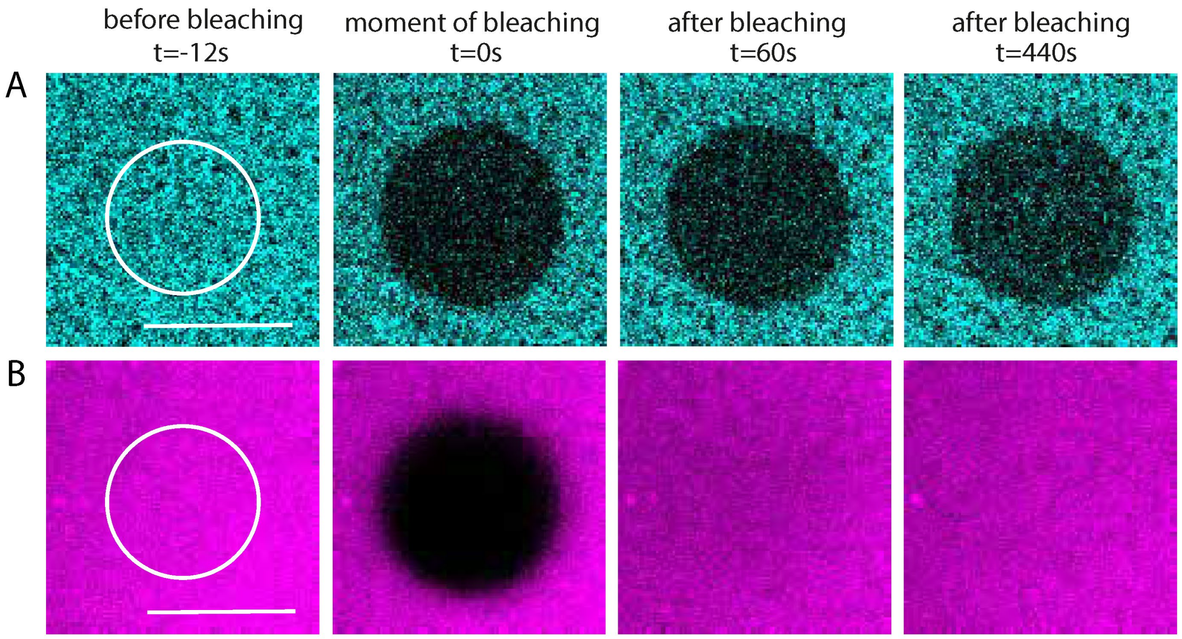


**Figure 2 – Figure supplement 2. Fluorescence recovery after photobleaching experiments show that membrane-adhered fly septin layers are immobile and stable over time while the bilayer is fluid.** A solution of 500 nM septin hexamers containing 10 mol-% mGFP-labeled hexamers was incubated with glass-supported lipid bilayers containing PC, 20% PS, and 0.3% mol-% of fluorescently labeled lipids (rhodamine-PE). (A) The fluorescence of the septin meshwork does not recover after photobleaching a circular region with a radius $r$ = 5 µm (indicated by the white circle) at time $t$ = 0 for 1 s on an observation time scale of nearly 7 min, indicating negligible mobility and turnover. (B) The fluorescent signal of the rhodamine-PE lipids does recover fully upon photobleaching. The time-dependent fluorescence recovery reveals an average lipid diffusivity $D \approx 1.2$ $\mu m^2/s$, both with and without adsorbed septins, as determined by extracting the recovery half time $\tau_{1/2} = 4r2\gamma_D/D$ (where $\gamma_D = 0.88$) from a fit to the Soumpasis equation [2]. This diffusivity is characteristic of fluid lipid bilayers on solid supports [3]. Scale bar: 10 µm.

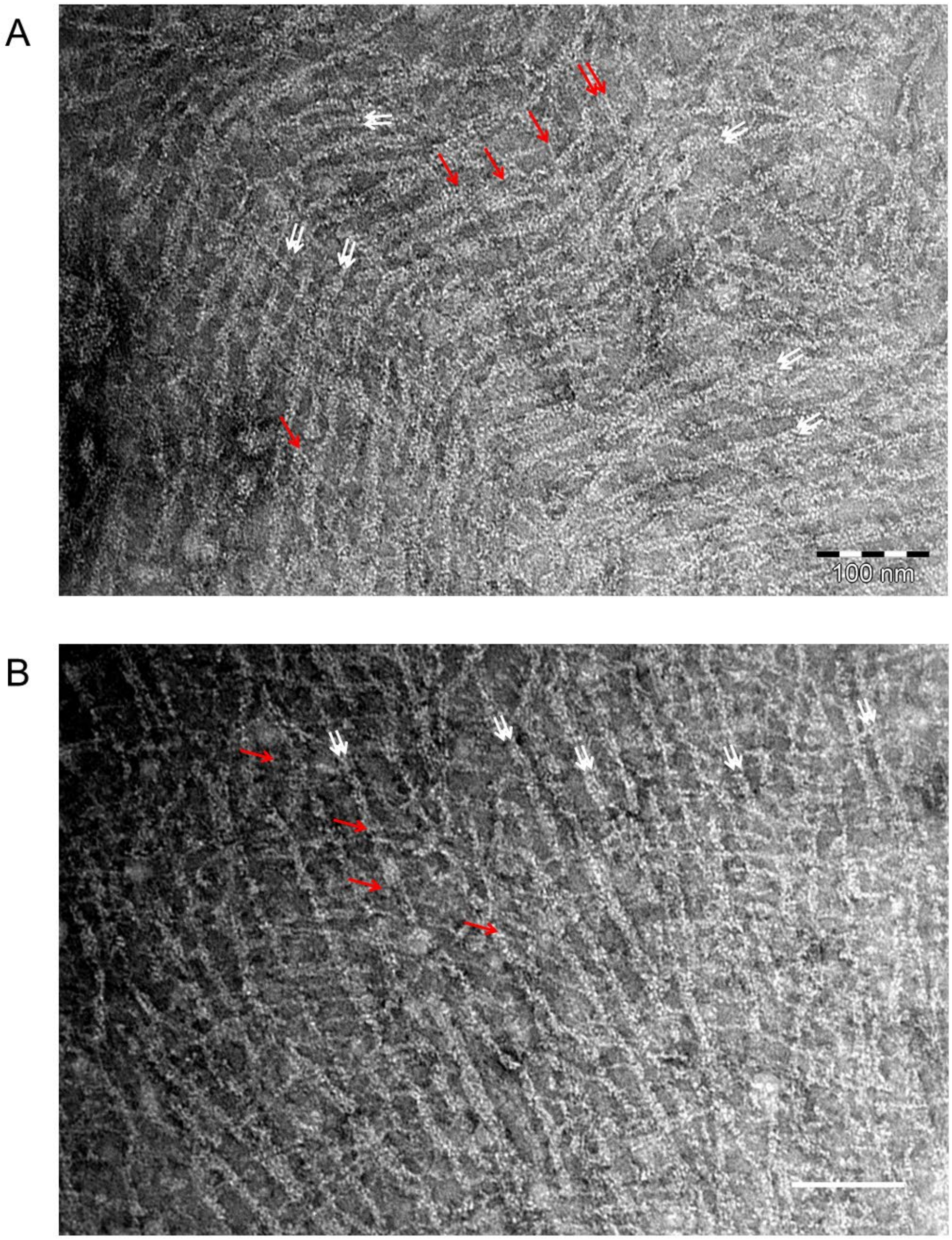


**Figure 3 – Figure supplement 1. Mammalian septin hexamers form paired filaments on lipid monolayers comprised of PC, 20 mol-% PS, and 5 mol-% PI(4,5)$P_2$.** Mammalian septin hexamers (comprised of mSept2, mSept6 and mSept7) incubated at 70 nM with a lipid monolayer containing 20% PS and 5% PIP2 form paired filaments connected by many cross-bridging filaments. The white arrows indicate examples of paired filaments (in areas where there are two obvious densities side by side and the filament as a whole looks thicker). Red arrows point at bridging filaments which seem to be thinner and run continuously across the paired filaments. It is unclear whether the bridging filaments are single filaments or (twisted, and therefore thinner) paired filaments. Scale bars: 100 nm.

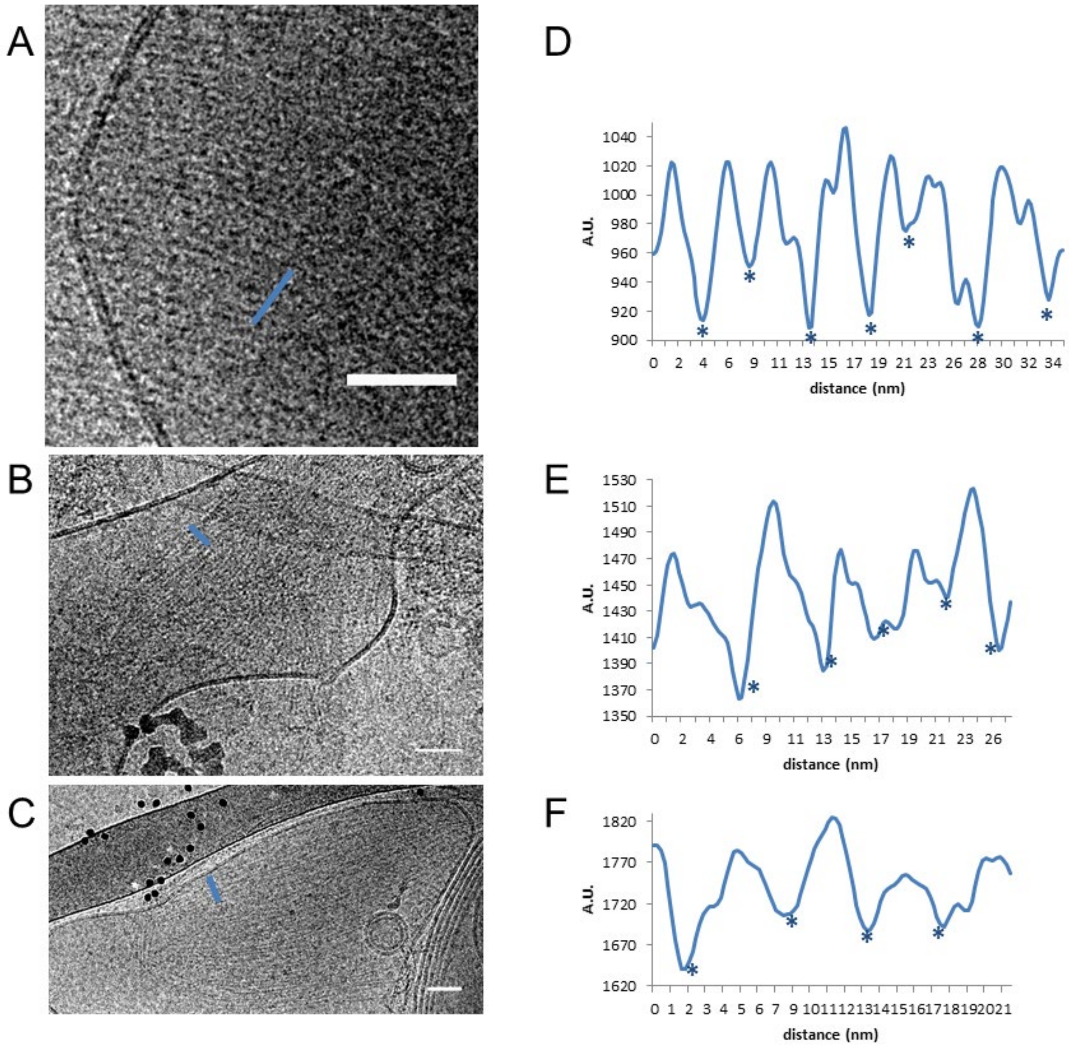


**Figure 4 – Figure supplement 1. Paired fly septin filaments on large unilamellar lipid vesicles exhibit an average spacing of ~5.7 nm.** (A, B, C) Three examples of cryoEM images, with (D, E, F) corresponding line profiles drawn along the blue lines. Stars denote approximate center positions of filaments. Scale bars: 50 nm.

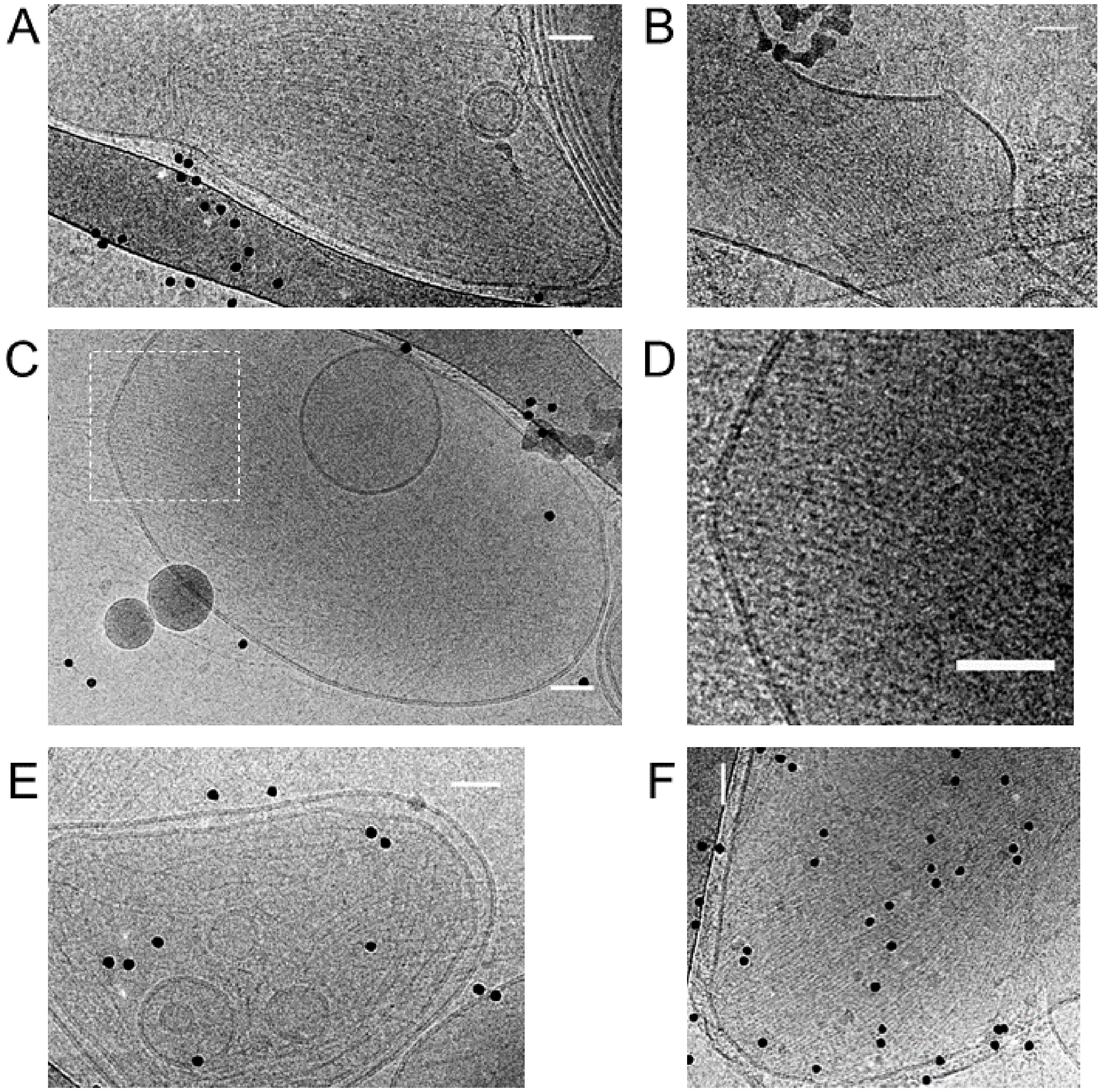


**Figure 4 – Figure supplement 2. Fly septin hexamers form paired and occasionally single filaments on large unilamellar lipid vesicles.** Gallery of CryoEM images of septins (300 nM) after a 30 minute incubation with vesicles composed of PC : PE : PI(4,5)$P_2$ at molar ratios of 83 : 11 : 6. PE was included based on prior cryoEM studies of protein binding to PI(4,5)$P_2$-containing membranes [4, 5]. Images are from different regions/samples, except for panel D, which shows a zoom-in of the boxed region in C. The images show septin filaments (linear structures) and vesicles of different sizes and shapes (circular objects). Most vesicles are covered with parallel sheets of tightly paired filament with narrow spacing (e.g. panel A), but some display rather spaced parallel filaments (*e.g.* panel E). Black dots are 10 nm diameter gold beads used as fiducials for cryo-tomography. Scale bars: 50 nm.

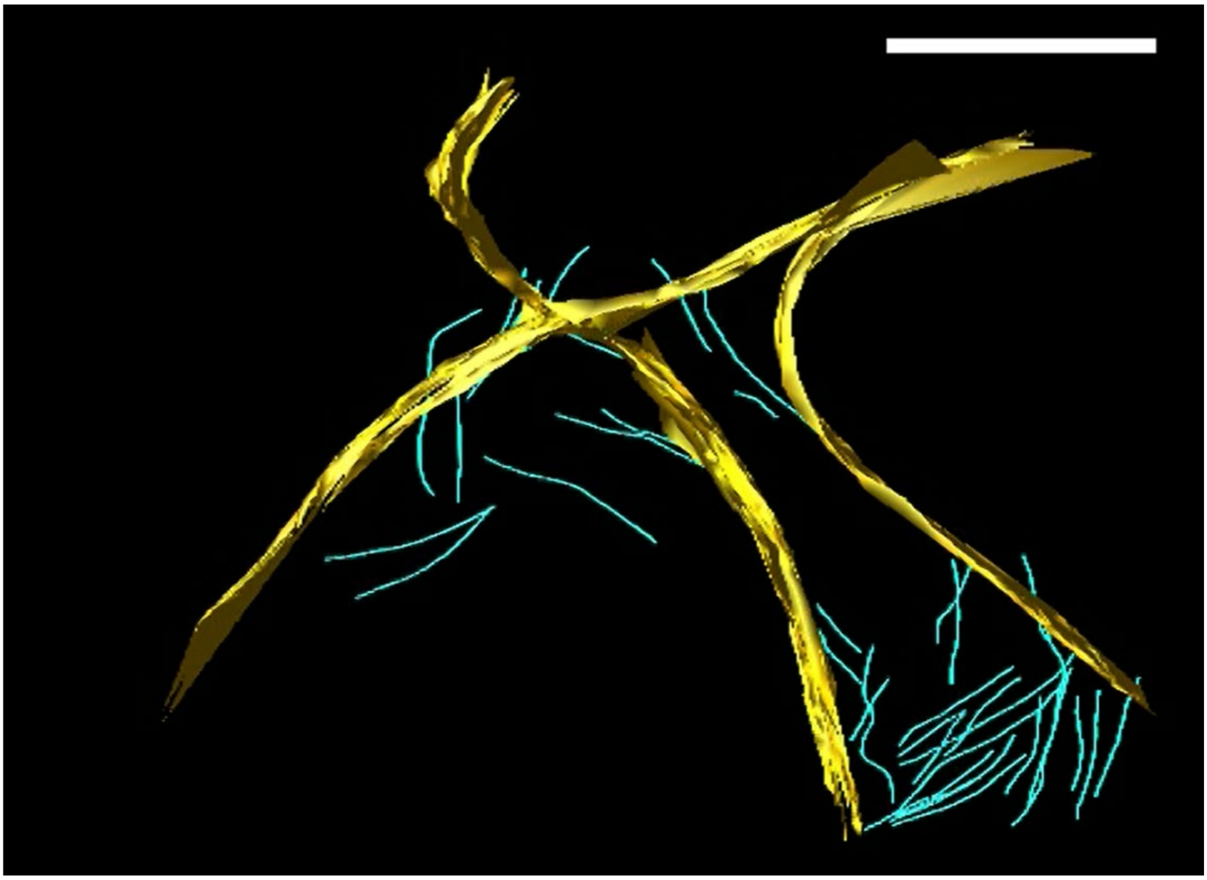

**Figure 4 – Video supplement 1 [**here: still image**].** 3D tomographic reconstruction of septins bound to lipid vesicles. The membrane is in yellow while septin filaments are segmented in blue. Scale bar: 850 nm. The membrane (though only partly visible because a part is perpendicular to the electron beam and therefore invisible because of the missing wedge in tomography) is distorted because of the bound septin filaments.

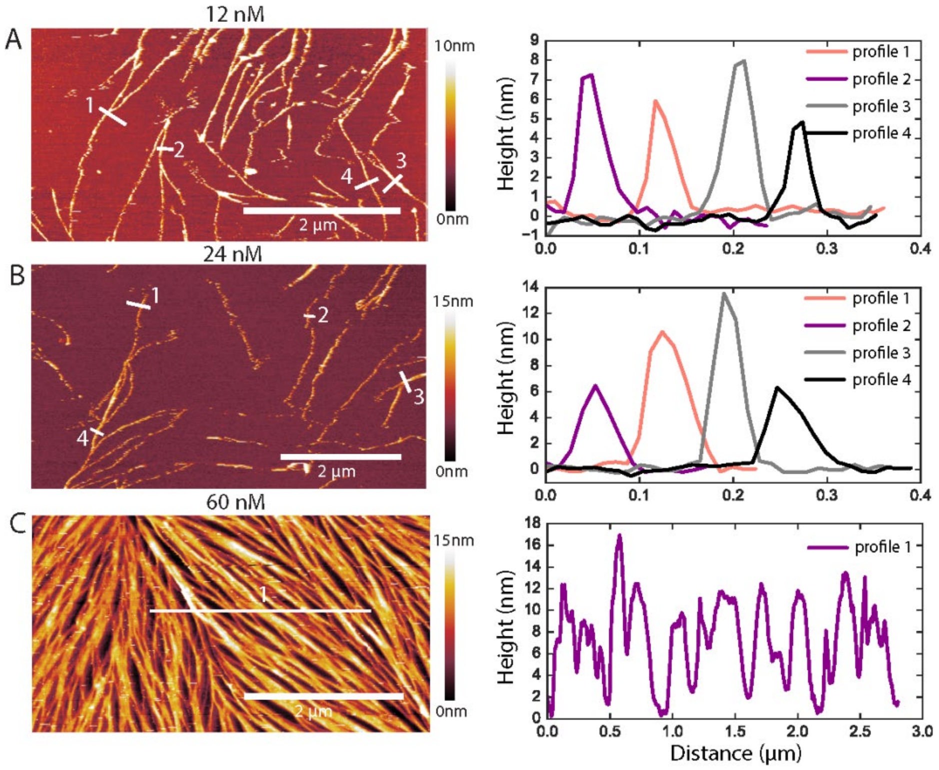


**Figure 5 – Figure supplement 1. Examples of AFM height profiles across filaments and bundles.** (Left) AFM topographic images of GTA-fixed fly septins at concentrations of 12 nM (A), 24 nM (B), and 60 nM (C). The images were cropped from the images presented in Figure 5B, and color bars on the right show the height scale. (Right) Height profiles were determined along the numbered white lines in each corresponding image on the left. In A and B, they run perpendicularly to the filament/bundle axes, as done for height and width analysis in Figure 5C. In C, a horizontal line across many filaments/bundles was chosen to appreciate their rather dense packing: note that the AFM tip, owing to its finite size, is unable to reach 0 nm height (*i.e.* the SLB surface) between some tightly spaced filament/bundles.

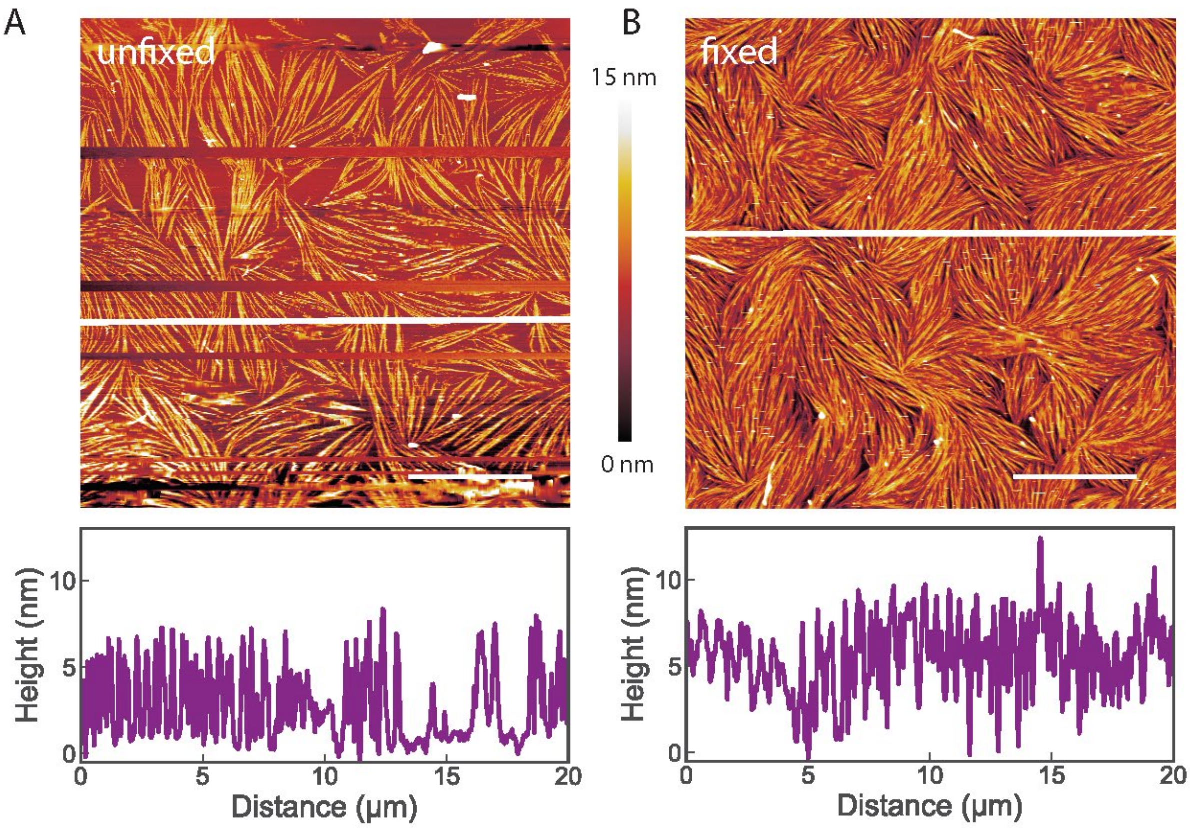


**Figure 5 – Figure supplement 2. AFM images show that glutaraldehyde (GTA) fixation does not change the morphology of fly septins adsorbed to supported lipid bilayers.** Septin hexamers were incubated at 60 nM on supported lipid bilayers made from 80 mol-% PC and 20 mol-% PS. The bundles and their liquid-crystalline organisation are preserved. (A) Image of unfixed septin filaments. (B) Image of septin filaments fixed by GTA treatment. Scale bar: 5 µm; color bar between images shows the height scale. The graphs below the images present height profiles along the solid white line depicted on each image. Note that the two samples were prepared independently, and that differences in surface density are likely due to slight variations in the sample preparation conditions. Occasional lines appearing empty (in A) correspond to phases when the AFM tip lost contact with the surface, as imaging without GTA required minimal forces to avoid perturbation of the septin organisation.

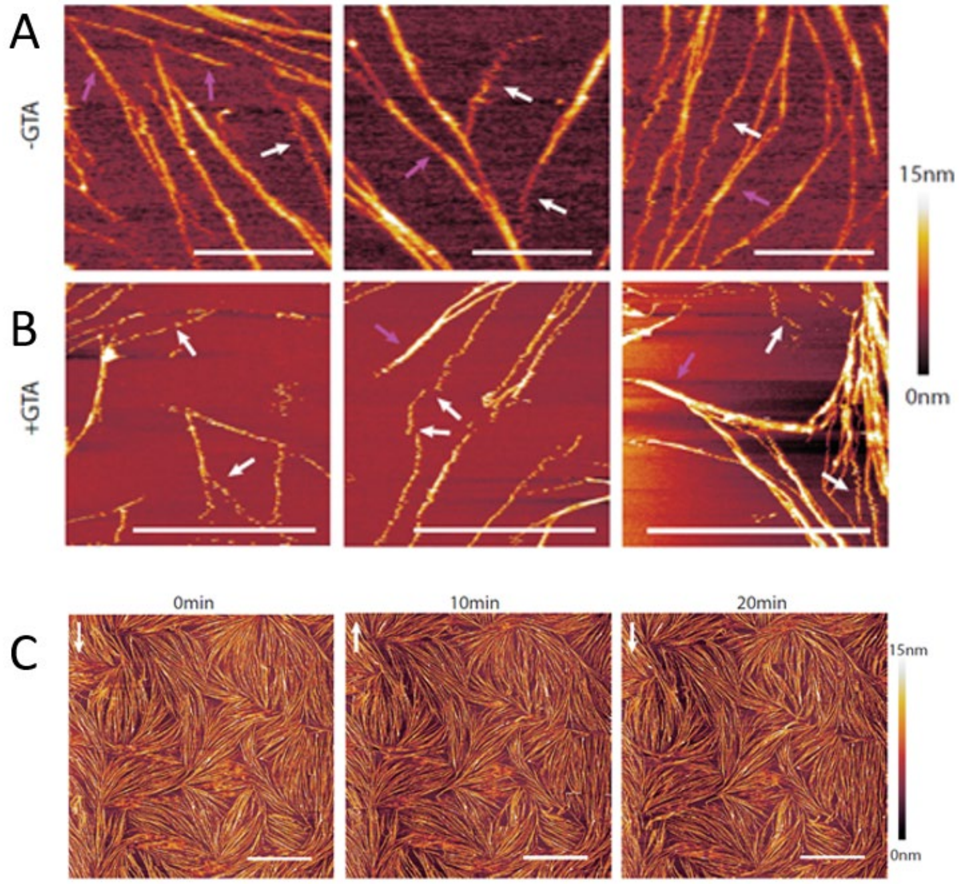


**Figure 5 – Figure supplement 3. AFM imaging shows that fly septin bundles and dense septin arrays are more mechanically stable than isolated septin filaments.** (A) Three representative images taken at different surface areas in unfixed conditions (-GTA). (B) Three representative images taken at different surface areas after fixation with 1% GTA (+GTA). Septin concentration was 60 nM in A, and 24 nM in B, on supported lipid bilayers made from 80 mol-% PC and 20 mol-% PS. White arrows in A and B show examples of single septin filaments that appear perturbed and mechanically unstable, as indicated by a 'zigzag' shape, most likely due to the AFM tip moving the filament back and forth across the membrane. Pink arrows show (typically thicker) stable filaments or bundles. (C) Dense arrays of septins (60 nM) are stable, as demonstrated by three consecutive scans of the same region, each scan taking 10 min ($t$ = 0 min indicates the initiation of the first scan). White vertical arrows show the direction of the slow scan axis. Scale bars: 5 µm; color bars on the right show the height scales.

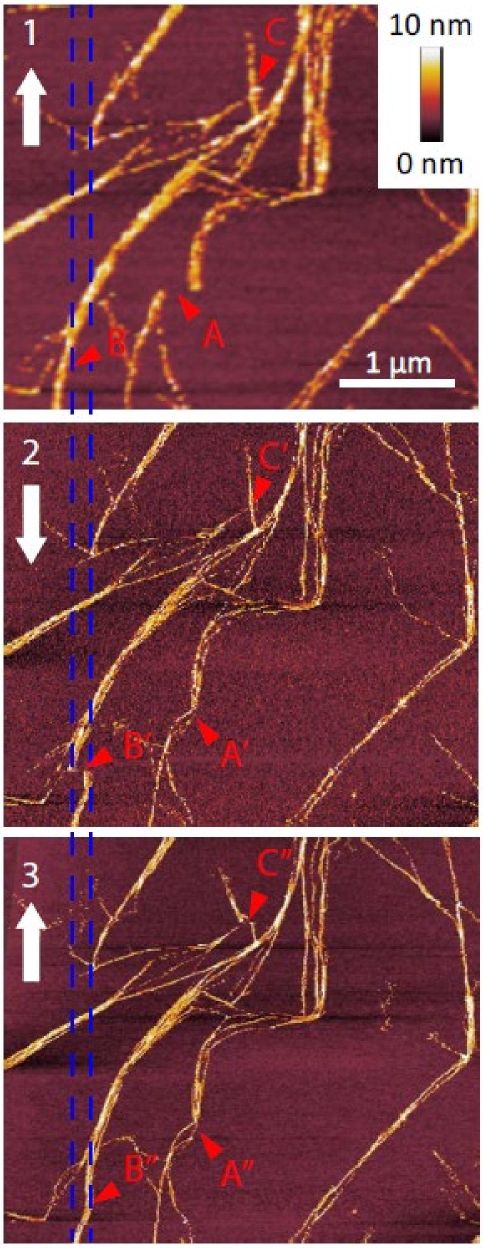


**Figure 5 – Figure supplement 4. Fly septin filament bundles can be displaced along the membrane.** Three consecutive atomic force micrographs (1, 2 and 3) of the same surface area are shown of GTA-fixed septin filaments (60 nM) on supported lipid bilayers made from 80 mol-% PC and 20 mol-% PS. The alternating direction of the slow-scan axis is indicated (white arrows). Most of the septin filaments remain in place, but some fibrils are occasionally displaced along the horizontal fast-scan axis owing to lateral forces exerted by the AFM tip. Arrow sequence B-B'-B'' shows such a displacement event leading to a permanent local kink and filament relocation (B''). The disruption of the filament (B') in micrograph 2 is most likely apparent, owing to the line-by-line scanning nature of AFM image acquisition. The spacing between the two blue vertical lines, corresponding to the displacement of the filament bundle in B', is 170 nm. Arrow sequences A-A'-A'' and C-C'-C'' highlight additional examples of displaced fibrils. Images 2 and 3 were stretched/compressed along the vertical (slow scan) axis and skewed along the horizontal (fast scan) axis to correct for drifts during imaging and align these images with image 1.

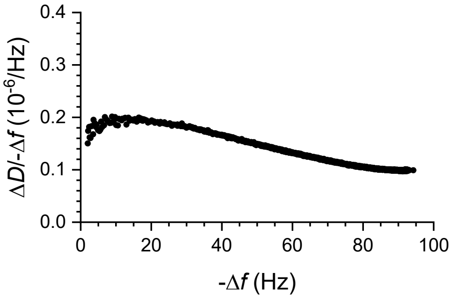


**Figure 7 – Figure supplement 1. Fly septin filament and bundles are sensed as soft relative to membrane-bound globular proteins by QCM-D.** The plot shows $\Delta D/-\Delta f$ ratios (a measure of elastic compliance, or softness) as a function of $-\Delta f$ (a measure of coverage) for septins incubated at 60 nM on an SLB made from 80% DOPC and 20% DOPS (data taken from Figure 7B). $\Delta D/-\Delta f$ ratios are relatively high, consistent with flexible hinges linking the protein to the membrane and/or inter-connecting protein domains.
